# Cortical Representations of Speech in a Multi-talker Auditory Scene

**DOI:** 10.1101/124750

**Authors:** Krishna C. Puvvada, Jonathan Z. Simon

## Abstract

The ability to parse a complex auditory scene into perceptual objects is facilitated by a hierarchical auditory system. Successive stages in the hierarchy transform an auditory scene of multiple overlapping sources, from peripheral tonotopically-based representations in the auditory nerve, into perceptually distinct auditory-objects based representation in auditory cortex. Here, using magnetoencephalography (MEG) recordings from human subjects, both men and women, we investigate how a complex acoustic scene consisting of multiple speech sources is represented in distinct hierarchical stages of auditory cortex. Using systems-theoretic methods of stimulus reconstruction, we show that the primary-like areas in auditory cortex contain dominantly spectro-temporal based representations of the entire auditory scene. Here, both attended and ignored speech streams are represented with almost equal fidelity, and a global representation of the full auditory scene with all its streams is a better candidate neural representation than that of individual streams being represented separately. In contrast, we also show that higher order auditory cortical areas represent the attended stream separately, and with significantly higher fidelity, than unattended streams. Furthermore, the unattended background streams are more faithfully represented as a single unsegregated background object rather than as separated objects. Taken together, these findings demonstrate the progression of the representations and processing of a complex acoustic scene up through the hierarchy of human auditory cortex.

**Significance Statement:** Using magnetoencephalography (MEG) recordings from human listeners in a simulated cocktail party environment, we investigate how a complex acoustic scene consisting of multiple speech sources is represented in separate hierarchical stages of auditory cortex. We show that the primary-like areas in auditory cortex use a dominantly spectro-temporal based representation of the entire auditory scene, with both attended and ignored speech streams represented with almost equal fidelity. In contrast, we show that higher order auditory cortical areas represent an attended speech stream separately from, and with significantly higher fidelity than, unattended speech streams. Furthermore, the unattended background streams are represented as a single undivided background object rather than as distinct background objects.

## Introduction

Individual sounds originating from multiple sources in a complex auditory scene mix linearly and irreversibly before they enter the ear, yet are perceived as distinct objects by the listener (Cherry, 1953; Bregman, 1994; McDermott, 2009). The separation, or rather individual re-creation, of such linearly mixed original sound sources is a mathematically ill-posed question, yet the brain nevertheless routinely performs this task with ease. The neural mechanisms by which this perceptual ‘un-mixing’ of sounds occur, the collective cortical representations of the auditory scene and its constituents, and the role of attention in both, are key problems in contemporary auditory neuroscience.

It is known that auditory processing in primate cortex is hierarchical (Davis and Johnsrude, 2003; Hickok and Poeppel, 2007; Rauschecker and Scott, 2009; Okada et al., 2010; Peelle et al., 2010; Overath et al., 2015) with subcortical areas projecting onto the core areas of auditory cortex, and from there, on to belt, parabelt and additional auditory areas (Kaas and Hackett, 2000). Sound entering the ear reaches different anatomical/functional areas of auditory cortex with different latencies (Recanzone et al., 2000; Nourski et al., 2014). Due to this serial component of auditory processing, the hierarchy of processing can be described by both anatomy and latency, of which the latter may be exploited using the high temporal fidelity of non-invasive magnetoencephalography (MEG) neural recordings.

In selective listening experiments using natural speech and MEG, the two major neural responses known to track the speech envelope are the M50_TRF_ and M100_TRF_, with respective latencies of 30 – 80 ms and 80 – 150 ms, of which the dominant neural sources are, respectively, Heschl’s gyrus (HG) and Planum temporale (PT) (Steinschneider et al., 2011; Ding and Simon, 2012b). Posteromedial HG is the site of core auditory cortex; PT contains both belt and parabelt auditory areas (here collectively referred to as higher-order areas) (Griffiths and Warren, 2002; Sweet et al., 2005). Hence the earlier neural responses are dominated by core auditory cortex, and the later are dominated by higher-order areas. To better understand the neural mechanisms of auditory scene analysis, it is essential to understand how the cortical representations of a complex auditory scene change from the core to the higher order auditory areas.

One topic of interest is whether the brain maintains distinct neural representations for each unattended source (in addition to the representation of the attended source), or if all unattended sources are represented collectively as a single monolithic background object. A common paradigm used to investigate the neural mechanisms underlying auditory scene analysis employs a pair of speech streams, of which one is attended, which then leaves the other speech stream remaining as the background (Kerlin et al., 2010; Ding and Simon, 2012b; Mesgarani and Chang, 2012; Power et al., 2012; Zion Golumbic et al., 2013b; O’Sullivan et al., 2015). This results in a limitation, which cannot address the question of distinct vs. collective neural representations for unattended sources. This touches on the long-standing debate of whether auditory object segregation is pre-attentive or it is actively influenced by attention (Carlyon, 2004; Sussman et al., 2005; Shinn-Cunningham, 2008; Shamma et al., 2011). Evidence for segregated neural representations of background streams would support the former, whereas a lack of segregated background objects would support the latter.

To address these issues, we use MEG to investigate a variety of potential cortical representations of the elements of a multi-talker auditory scene. We test two major hypotheses: that the dominant representation in core auditory cortex is of the physical acoustics, not of separated auditory objects; and that once object-based representations emerge in higher order auditory areas, the unattended contributions to the auditory scene are represented collectively as a single background object. The methodological approach employs the linear systems methods of stimulus prediction and MEG response reconstruction (Lalor et al., 2009; Mesgarani et al., 2009; Ding and Simon, 2012b; Mesgarani and Chang, 2012; Pasley et al., 2012; Di Liberto et al., 2015).

## Materials & Methods

### Subjects & Experimental Design

Nine normal-hearing, young adults (6 Female) participated in the experiment. All subjects were paid for their participation. The experimental procedures were approved by the University of Maryland Institutional Review Board. Subjects listened to a mixture of three speech segments spoken by, respectively, a male adult, female adult and a child speaker. The three speech segments were mixed into a single audio channel with equal perceptual loudness. All three speech segments were taken from public domain narration of Grimms’ Fairy Tales by Jacob & Wilhelm Grimm (https://librivox.org/fairy-tales-by-the-brothers-grimm/). Periods of silence longer than 300 ms were replaced by a shorter gap whose duration was chosen randomly between 200 ms and 300 ms. The audio signal was low-pass filtered with cut-off at 4 kHz. In first of three conditions, the subjects were asked to attend to the child speaker, while ignoring the other two (i.e., child speaker as target, with male and female adult speakers as background). In condition two, during which the same mixture was played as in condition one, the subjects were instead asked to attend to the male adult speaker (with female adult and child speakers as background). Similarly, in condition three, the target was switched to the female adult speaker. Each condition was repeated three times successively, producing three trials per condition. The presentation order of the three conditions was counterbalanced across subjects. Each trial was of 220 s duration, divided into two 110 s partitions, to reduce listener fatigue. To help participants attend to the correct speaker, the first 30 s of each partition was replaced by the clean recording of the target speaker alone, followed by a 5 s upward linear ramp of the background speakers. Recordings of this first 35 s of each segment were not included in any analysis. To further encourage the subjects to attend to the correct speaker, a target-word was set before each trial and the subjects were asked to count the number of occurrences of the target-word in the speech of the attended speaker. Additionally, after each condition, the subject was asked to recount a short summary of the attended narrative. The subjects were required to close their eyes while listening. Before the main experiment, 100 repetitions of a 500-Hz tone pip were presented to each subject to elicit the M100 response, a reliable auditory response occurring ~100 ms after the onset of a tone pip. This data was used check whether any potential subjects gave abnormal auditory responses, but no subjects were excluded based on this criterion.

### Data recording and pre-processing

MEG recordings were conducted using a 160-channel whole-head system (Kanazawa Institute of Technology, Kanazawa, Japan). Its detection coils are arranged in a uniform array on a helmet-shaped surface of the bottom of the dewar, with ~25 mm between the centers of two adjacent 15.5-mm-diameter coils. Sensors are configured as first-order axial gradiometers with a baseline of 50 mm; their field sensitivities are 5 fT/√Hz or better in the white noise region. Subjects lay horizontally in a dimly lit magnetically shielded room (Yokogawa Electric Corporation). Responses were recorded with a sampling rate of 1 kHz with an online 200-Hz low-pass filter and 60 Hz notch filter. Three reference magnetic sensors and three vibrational sensors were used to measure the environmental magnetic field and vibrations. The reference sensor recordings were utilized to reduce environmental noise from the MEG recordings using the Time-Shift PCA method (de Cheveigne and Simon, 2007). Additionally, MEG recordings were decomposed into virtual sensors/components using denoising source separation (DSS) (Särelä and Valpola, 2005; de Cheveigne and Simon, 2008; de Cheveigne and Parra, 2014), a blind source separation method that enhances neural activity consistent over trials. Specifically, DSS decomposes the multichannel MEG recording into temporally uncorrelated components, where each component is determined by maximizing its trial-to-trial reliability, measured by the correlation between the responses to the same stimulus in different trials. To reduce the computational complexity, for all further analysis the 157 MEG sensors were reduced, using DSS, to 4 components in each hemisphere. Also, both stimulus envelope and MEG responses were band pass filtered between 1 – 8 Hz (delta and theta bands), which correspond to the slow temporal modulations in speech (Ding and Simon, 2012a, b).

### Neural Model Terminology and Notation

As specified in the stimulus description, in each condition the subject attends to one among the three speech streams. Neural models of speech stream processing can be compared by contrasting the predicted envelope reconstructions of the different models. The envelope of attended speech stream is referred to as the ‘foreground’ and the envelope of each of the two unattended speech streams is referred to as the ‘individual background’. In contrast, the envelope of the entire unattended part of the stimulus, comprising *both* unattended speech streams, is referred to as the ‘combined background’. The envelope of entire acoustic stimulus or auditory scene, comprising of all the three speech streams is referred to as the ‘acoustic scene’. Thus, if *S*_a_, *S*_b_, *S*_c_ are three speech stimuli, *Env* (*S*_a_ + *S*_b_ + *S*_c_) is the acoustic scene. In contrast, the sum of envelopes of three speech streams, *Env* (*S*_*a*_)+ *Env* (*S*_*b*_)+ *En* (*S*_*c*_), is referred to as the ‘sum of streams’, and the two are not mathematically equal: even though both are functions of the same stimuli, they differ due to the non-linear nature of a signal envelope (the linear correlation between the acoustic scene and the sum of streams is typically ~0.75). Combination (unsegregated) envelopes, whether of the entire acoustic scene or the background only, can be used to test neural models that do not perform stream segregation. Sums of individual stream envelopes, whether of all streams or just the background streams, can be used to test neural models that process the (segregated) streams in parallel, given that neurally generated magnetic fields add in linear superposition.

Neural responses with latencies less than ~85 ms (typically originating from core auditory areas) are referred to here as ‘early neural responses’ and responses with latencies more than ~85 ms (typically from higher-order auditory areas) (Ahveninen et al., 2011; Okamoto et al., 2011; Steinschneider et al., 2011) are referred to as ‘late neural responses’.

The next two sections describe models of the neural encoding of stimuli into responses, followed by models of the decoding of stimuli from neural responses. Encoding models are presented here first because of their ease of description over decoding models, but in Results the decoding analysis is presented first, since it is the decoding results that inform the new model of encoding.

### Temporal Response Function

In an auditory scene with a single talker, the relation between MEG neural response and the presented speech stimuli can be modeled using a linear temporal response function (TRF) as

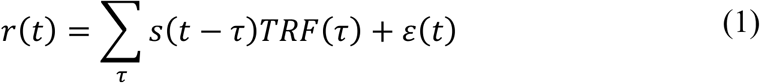

where *t* = 0,1, …, *T* is time, *r t* is the response from any individual sensor or DSS component, *s* (*t*) is the stimulus envelope in decibels, *TRF* (*t*) is the TRF itself, and (*t*) is residual response waveform not explained by the TRF model (Ding and Simon, 2012a). The envelope is extracted by averaging the auditory spectrogram, (Chi et al., 2005) along the spectral dimension. The TRF is estimated using boosting with 10-fold cross-validation (David et al., 2007). In case of single speech stimuli, the TRF is typically characterized by a positive peak between 30 ms and 80 ms and a negative peak between 90 ms and 130 ms, referred to as M50_TRF_ and M100_TRF_ respectively (Ding and Simon, 2012b) (positivity/negativity of the magnetic field is by convention defined to agree with the corresponding electroencephalography[EEG] peaks). Success/accuracy of the linear model is evaluated by how well it predicts neural responses, as measured by the proportion of the variance explained: the square of the Pearson correlation coefficient between the MEG measurement and the TRF model prediction.

In the case of more than one speaker, the MEG neural response, *r* (*t*) can be modeled as the sum of the responses to the individual acoustic sources (Ding and Simon, 2012b; Zion Golumbic et al., 2013b), referred to here as the ‘Summation model’. For example, with three speech streams, the neural response would be modeled as

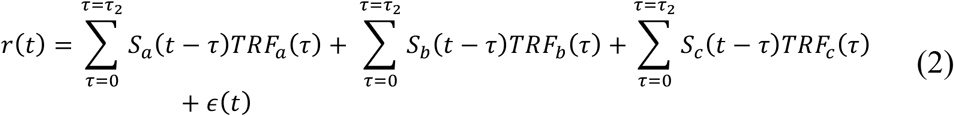

where *S*_*a*_ (*t*), *S*_*S*_(*t*) and *S*_*c*_(*t*) are the envelopes of the three speech streams, and *TRF*_*a*_ (*t*), *TRF*_*b*_ (*t*) and *TRF*_*c*_(*t*) are the TRFs corresponding to each stream. τ_2_represents the length of TRF. All TRFs in the Summation model are estimated simultaneously.

In addition to the existing summation model, we propose a new encoding-model referred to as the ‘Early-late model’, which allows one to incorporate the hypothesis that the early neural responses typically represent the entire acoustic scene, but that the later neural responses differentially represent the separated foreground and background.

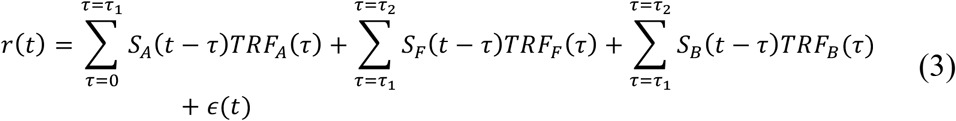

where *S*_A_(*t*) is the (entire) acoustic scene, *S*_F_(*t*) is the envelope of attended (foreground) speech stream, and *S*_C_(*t*) is the combined background (i.e., envelope of everything other than attended speech stream in the auditory scene), and *TRF*_A_ (*t*), *TRF*_F_ (*t*), and *TRF*_B_(*t*) are the corresponding TRFs. *τ*_1_, *τ*_2_ represent the boundary values of the integration windows for early and late neural responses respectively, with 0 < *τ*_1_ < *τ*_2_.

The explanatory power of different models, such as the Summation and Early-late models, can be ranked by comparing the accuracy of their response predictions (illustrated in Figure 1A). The models differ in terms of number of free parameters, with the Early-late model having fewer parameters than the Summation model. Hence, any improved performance observed in the proposed Early-late model over the Summation model is correspondingly more likely due to a better model fit, since it has less freedom to fit the data (though the converse would not hold).

**Figure 1:**
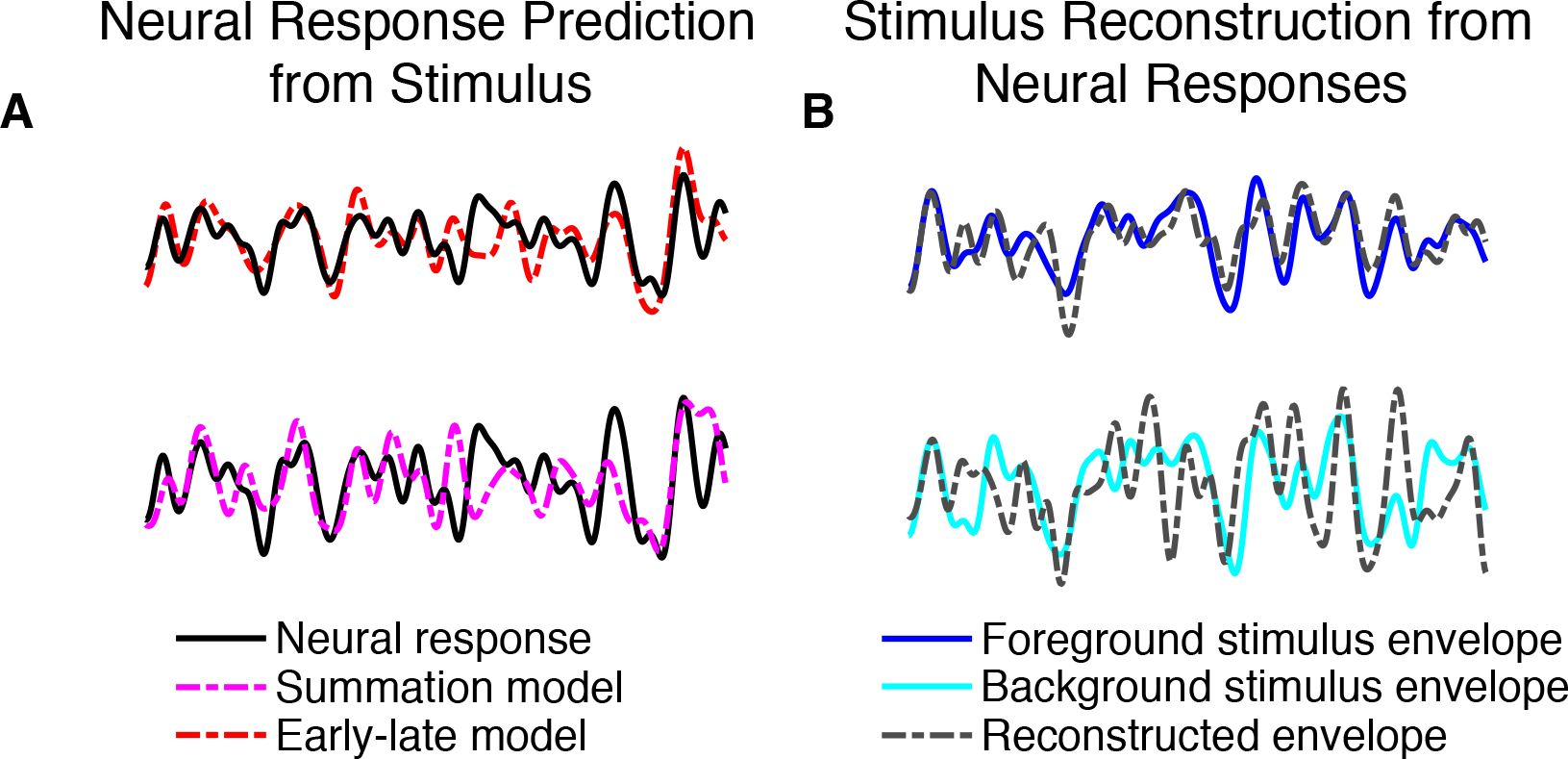
Illustrations of outcomes comparing competing encoding- and decoding-based neural representations of the auditory scene and its constituents. All examples are grand averages across subjects (3 seconds duration). **A.** Comparing competing models of *encoding* to neural responses. In both the top and bottom examples, an experimentally measured MEG response (black) is compared to the neural response predictions made by competing proposed models. In the top example, the neural response prediction (red) is from the Early-late model; in the bottom example, the neural response prediction (magenta) is from the Summation model. The proposed Early-late model prediction shows higher correlation with the actual MEG neural response than Summation model. **B.** Comparing competing models of *decoding* to stimulus speech envelopes. In both the top and bottom examples, an acoustic speech stimulus envelope (blue/cyan) is compared to the model reconstruction of the respective envelope (gray). In the top example, the envelope reconstruction (blue) is of the foreground stimulus, based on late time responses; in the bottom example, the envelope reconstruction (cyan) is of the background stimulus, also based on late time responses. The foreground reconstruction shows higher correlation with the actual foreground envelope, compared to the background reconstruction with the actual background envelope.

### Decoding speech from neural responses

While the TRF/encoding analysis described in the previous section predicts neural response from the stimulus, decoding analysis reconstructs the stimulus based on the neural response. Thus, decoding analysis complements the TRF analysis (Mesgarani et al., 2009). Mathematically the envelope reconstruction/decoding operation can be formulated as

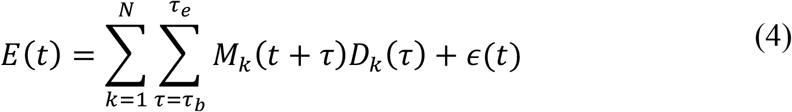

where *E* (*t*) is the reconstructed envelope, *M*_*k*_ (*t*) is the MEG recording (neural response) from sensor/component *k*, and *D*_*k*_ *t* is the linear decoder for sensor/component *k*. The times τ_*b*_ and τ_*e*_ denote the beginning and end times of the integration window. By appropriately choosing the values of τ_*b*_ and τ_*e*_, envelope reconstructions using neural responses from any desired time window can be compared. The decoder is estimated using boosting analogously to the TRF estimation in the previous section. In the single talker case the envelope is of that talker’s speech. In a multi-talker case, the envelope to be reconstructed might be the envelope of the speech of attended talker, or one of the background talkers, or of a mixture of any two or all three talkers, depending on the model under consideration. Chance-level reconstruction (i.e., the noise floor) from a particular neural response is estimated by reconstructing an unrelated stimulus envelope from that neural response. Figure 2 illustrates the distinction between reconstruction of stimulus envelope from early and late responses. The stimulus envelope at time point *t* can be reconstructed using neural responses from the dashed (early response) window or dotted (late response) window. (While it is true that the late responses to the stimulus at time point *t –*Δ*t* overlap with early responses to the stimulus at time point *t*, the decoder used to reconstruct the stimulus at time point *t* from early responses is only minimally affected by late responses to the stimulus at time point *t –*Δ*t* when the decoder is estimated by averaging over a long enough duration, e.g., tens of seconds). The cut-off time between early and late responses, *τ*_*boundry*_, was chosen to minimize the overlap between the M50_TRF_ and M100_TRF_ peaks, on a per subject basis, with a median value of 85 ms (range 70-100 ms in 5 ms increments); repeating the analysis using the single value of 85 ms for all subjects did not qualitatively change any conclusions. When decoding from early responses only, the time window of integration is from *τ*_*b*_ = 0 to *τ*_*e*_ = *τ*_*boundry*_. When decoding from late neural responses only, the time window of integration is from *τ*_*b*_ = *τ*_*boundry*_ to *τ*_*e*_ = 500 ms.

**Figure 2:**
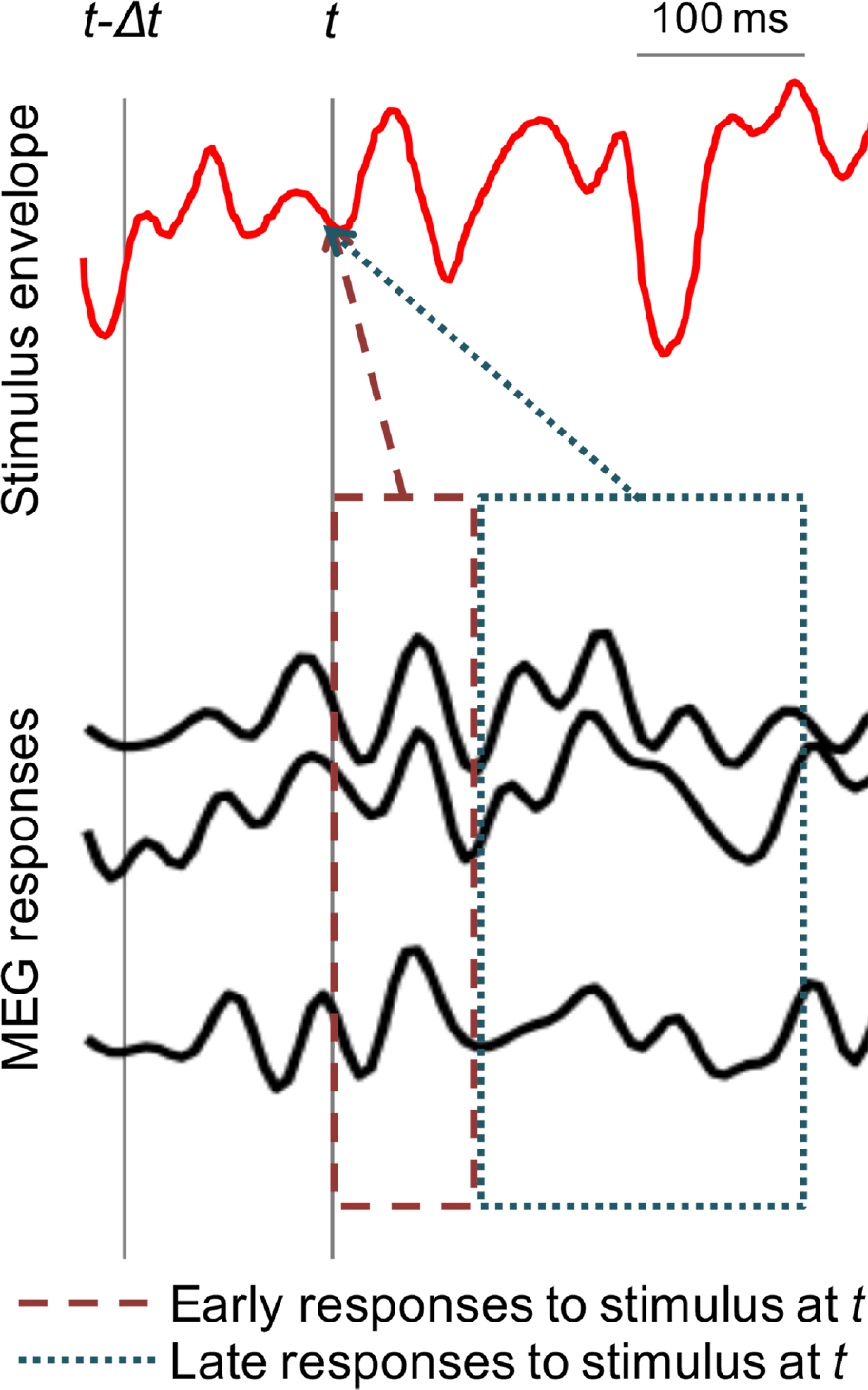
Early vs. late MEG neural responses to a continuous speech stimulus. A sample stimulus envelope and time-locked multi-channel MEG recordings are shown in red and black respectively. The two grey vertical lines indicate two arbitrary time points at *t – Δt* and *t*. The dashed and dotted boxes represent the early and late MEG neural responses to stimulus at time point *t* respectively. The reconstruction of the stimulus envelope at time *t* can be based on either early or late neural responses, and the separate reconstructions can be compared against each other.

The robustness of different representations, such as of Foreground vs. Background, can be compared by examining the accuracy of their respective stimulus envelope reconstructions (illustrated in Figure 1, right).

### Statistics

All statistical comparisons reported here are two-tailed permutation tests with *N*=1,000,000 random permutations (within subject). Due to the value of N selected, the smallest accurate *p* value that can be reported is 2×1/*N* (= 2×10^-6^; the factor of 2 arises from the two-tailed test) and any *p* value smaller than 2/*N* is reported as *p <* 2×10^-6^. The statistical comparison between foreground and individual backgrounds requires special mention, since each listening condition has one foreground but two individual backgrounds. From the perspective of both behavior and task, both the individual backgrounds are interchangeable. Hence, when comparing reconstruction accuracy of foreground vs. individual background the average reconstruction accuracy of the two individual backgrounds is used. Finally, Bayes factor analysis is used, when appropriate, to evaluate evidence in favor of null hypothesis, since conventional hypothesis testing is not suitable for such purposes. Briefly, Bayes factor analysis calculates the *posterior odds* i.e., the ratio of *P*(*H*_0_|*observations*) to *P*(*H*_1_|*observations*), where *H*_0_ and *H*_1_ are the null and alternate hypotheses respectively.

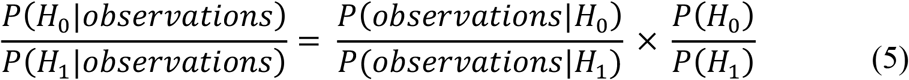

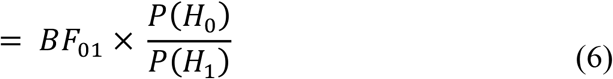

The ratio of *P*(*observations*|*H*_0_) and *P*(*observations*|*H*_1_) is denoted as the Bayes factor, BF_01_. Then, under the assumption of equal priors (*P*(*H*_0_) *= P*(*H*_1_)), the posterior odds reduces to BF_01_. A BF_01_ value of 10 indicates that the data is ten times more likely to occur under the null hypothesis than the alternate hypothesis; conversely, a BF_01_ value of 0.1 indicates that the data is 10 times more likely to occur under the alternate hypothesis than the null hypothesis. Conventionally, a BF_01_ value between 3 and 10 is considered as moderate evidence in favor of the null hypothesis, and a value between 10 and 30 is considered strong evidence; conversely, a BF_01_ value between 1/3 & 1/10 (respectively 1/10 & 1/30) is considered moderate (respectively strong) evidence for the alternate hypothesis (for more details we refer the reader to Rouder et al. (2009)).

## Results

### Stimulus reconstruction from early neural responses

To investigate the neural representations of the attended vs. unattended speech streams associated with early auditory areas, i.e., from core auditory cortex, (Nourski et al., 2014), the temporal envelope of attended (foreground) and unattended speech streams (individual backgrounds) were reconstructed using decoders optimized individually for each speech stream. All reconstructions performed significantly better than chance level (foreground vs. noise, *p <* 2×10^-6^; individual background vs. noise, *p <* 2×10^-6^), indicating that all three speech streams are represented in early auditory cortex. Figure 3A shows reconstruction accuracy for foreground vs. individual backgrounds. A permutation test shows no significant difference between foreground and individual background (*p* = 0.21), indicating that there is no evidence of significant neural bias for the attended speech stream over the ignored speech stream, in early neural responses. In fact, Bayes Factor analysis (BF_01_ = 4.2) indicates moderate support in favor of the null hypothesis (Rouder et al., 2009), that early neural responses do not distinguish significantly between attended and ignored speech streams.

**Figure 3:**
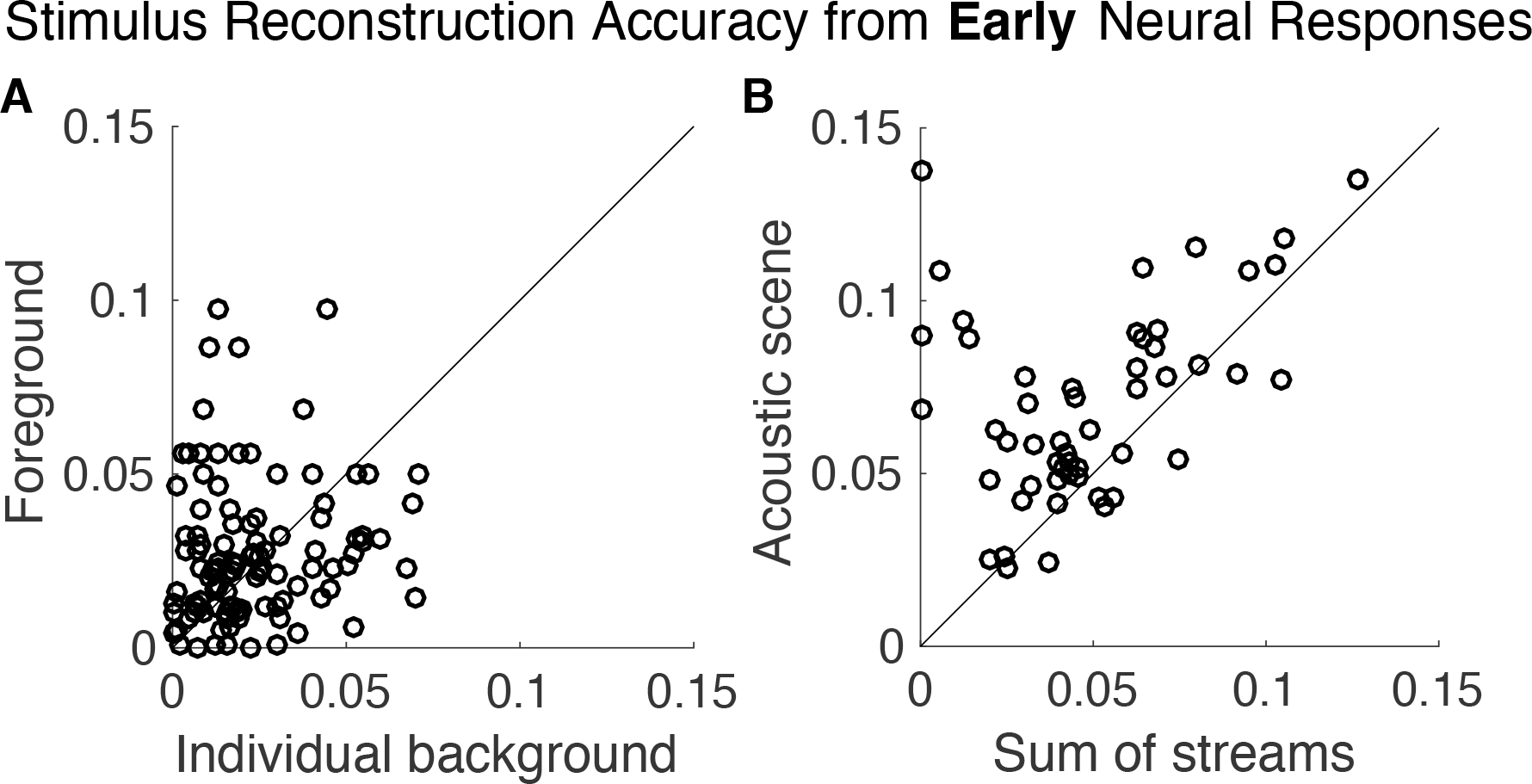
Stimulus envelope reconstruction accuracy using *early* neural responses. **A.** Scatter plot of reconstruction accuracy of the foreground vs. individual background envelopes. No significant difference was observed (*p* = 0.21), and therefore no preferential representation of the foreground speech over the individual background streams is revealed in early neural responses. Each data point corresponds to a distinct background and condition partition per subject (with two backgrounds sharing a common foreground). **B.** Scatter plot of reconstruction accuracy of the envelope of the entire acoustic scene vs. that of the sum of the envelopes of all three individual speech streams. The acoustic scene is reconstructed more accurately (visually, most of data points fall above the diagonal) as a whole than as the sum of individual components in early neural responses (*p <* 2 × 10^-6^). Each data point corresponds to a distinct condition partition per subject. In both plots, reconstruction accuracy is measured by proportion of the variance explained: the square of the Pearson correlation coefficient between the actual and predicted envelopes.

To test the hypothesis that early auditory areas represent the auditory scene in terms of acoustics, rather than as individual auditory objects, we reconstructed the acoustic scene (the envelope of the sum of all three speech streams) and compared it against the reconstruction of the sum of streams (sum of reconstruction envelopes of each of the three individual speech streams). Separate decoders optimized individually were used to reconstruct the acoustic scene and the sum of streams. As can be seen in Figure 3B, the result shows that the acoustic scene is better reconstructed than the sum of streams (*p <* 2×10^-6^). This indicates that early auditory cortex is better described as processing the entire acoustic scene rather than processing the separate elements of the scene individually.

### Stimulus reconstruction from late neural responses

While the preceding results were based on early cortical processing, the following results are based on late auditory cortical processing (responses with latencies more than ~85 ms). Figure 4A shows the scatter plot of reconstruction accuracy for the foreground vs. individual background envelopes based on late responses. A paired permutation test shows that reconstruction accuracy for the foreground is significantly higher than the background (*p <* 2×10^-6^). Even though the individual backgrounds are not as reliably reconstructed as foreground, their reconstructions are nonetheless significantly better than chance level (*p <* 2×10^-6^).

In order to distinguish among possible neural representations of the background streams, we compared the reconstructability of the envelope of the entire background as a whole, with the reconstructability of the sum of the envelopes of the (two) backgrounds. If the background is represented as a single auditory object (i.e., “the background”), the reconstruction of the envelope of the entire background should be more faithful than the sum of envelopes of individual backgrounds. In contrast, if the background is represented as distinct auditory objects, each distinguished by its own envelope, the reconstruction of the sum of envelopes of the individual backgrounds should be more faithful. Figure 4B shows the scatter plot of reconstruction accuracy for the envelope of combined background vs. the sum of the envelopes of the individual background streams. Analysis shows that the envelope of the combined background is significantly better represented than the sum of the individual envelopes of the individual backgrounds (*p* = 0.012). As noted previously, the envelope of the combined background is actually strongly correlated with the sum of the envelopes of the individual backgrounds, meaning that finding a significant difference in their reconstruction accuracy is *a priori* unlikely, providing even more credence to the result.

**Figure 4:**
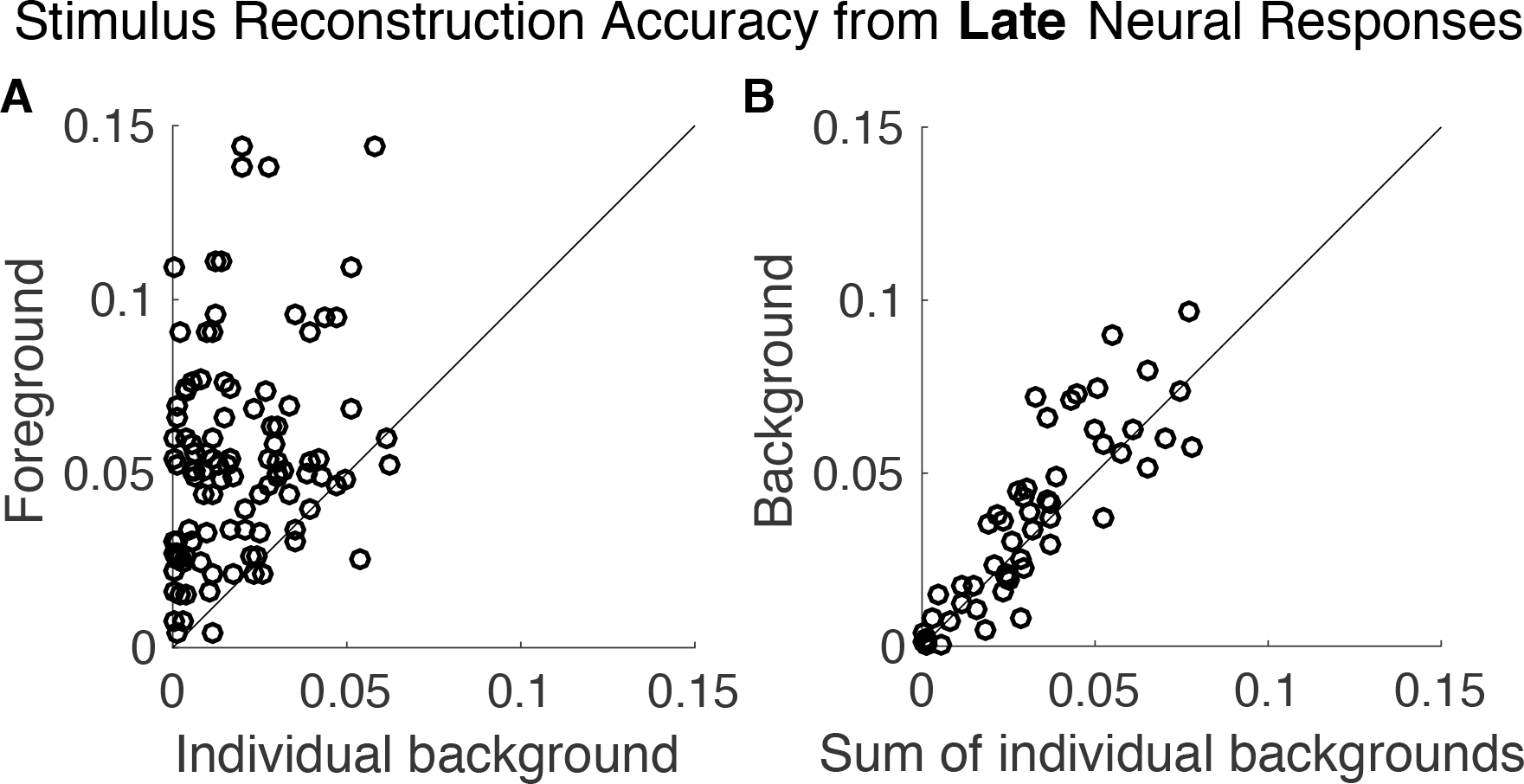
Stimulus envelope reconstruction accuracy using *late* neural responses. **A.** Scatter plot of accuracy between foreground vs. individual background envelope reconstructions demonstrates that the foreground is represented with dramatically better fidelity (visually, most of data points fall above the diagonal) than the background speech, in late neural responses (*p <* 2 × 10^-6^). Each data point corresponds to a distinct background and condition partition per subject (with two backgrounds sharing a common foreground). **B.** Scatter plot of the reconstruction accuracy of the envelope of the entire background vs. that of the sum of the envelopes of the two individual background speech streams. The background scene is reconstructed more accurately as a monolithic background than as separated individual background streams in late neural responses (*p* = 0.012). Each data point corresponds to a distinct condition partition per subject.

### Encoding analysis

Results above from envelope reconstruction suggest that while early neural responses represent the auditory scene in terms of the acoustics, the later neural responses represent the auditory scene in terms of a separated foreground and a single background stream. In order to further test this hypothesis, we use TRF-based encoding analysis to directly compare two different models of auditory scene representations. The two models compared are the standard Summation model (based on parallel representations of all speech streams; see Equation 2) and the new Early-late model (based on an early representation of the entire acoustic scene and late representations of separated foreground and background; see Equation 3). Figure 5 shows the response prediction accuracies for the two models. A permutation test shows that the accuracy of the Early-late model is considerably higher than that of the Summation model (*p <* 2×10^-6^). This indicates that a model in which early/core auditory cortex processes the entire acoustic scene but later/higher-order auditory cortex processes the foreground and background separately has more support than the previously employed model of parallel processing of separate streams throughout auditory cortex.

**Figure 5:**
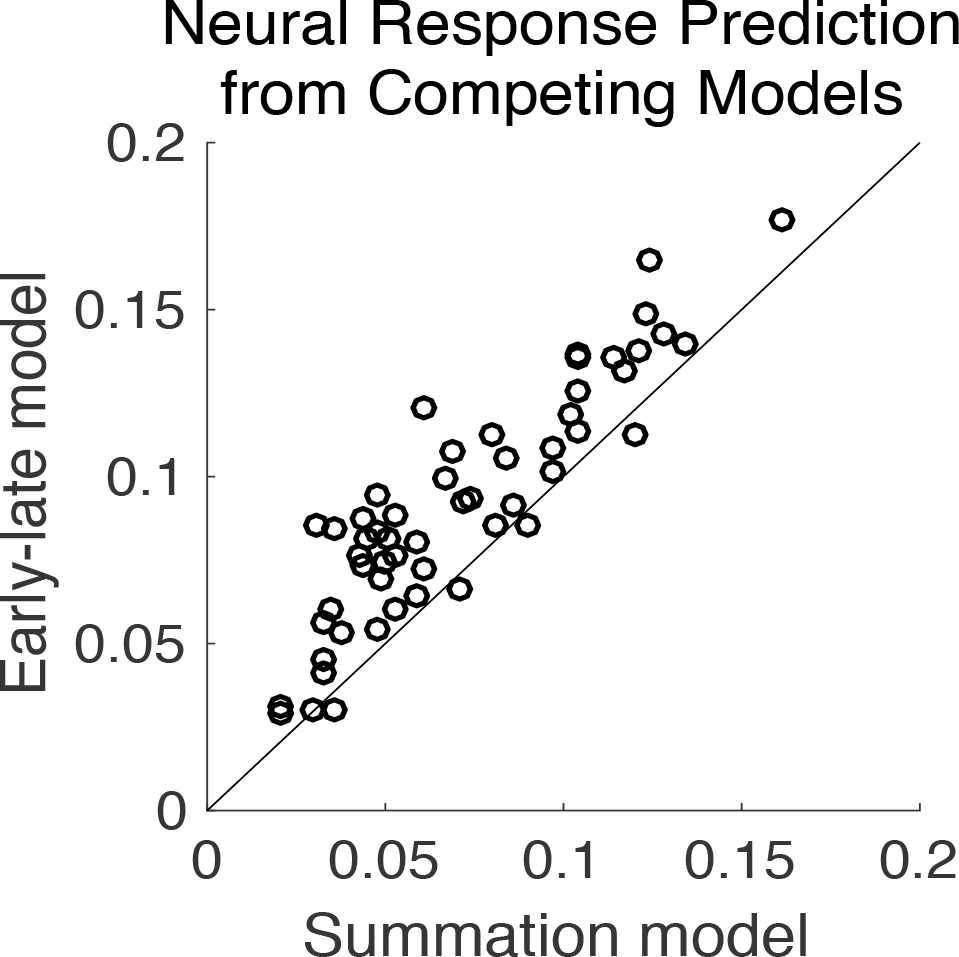
MEG response prediction accuracy. Scatter plot of the accuracy of predicted MEG neural response for the proposed Early-late model vs. the standard Summation model. The Early-late model predicts the MEG neural response dramatically better (visually, most of data points fall above the diagonal) than the Summation model (*p <* 2 × 10^-6^). The accuracy of predicted MEG neural responses is measured by proportion of the variance explained: the square of the Pearson correlation coefficient between the actual and predicted responses. Each data point corresponds to a distinct condition partition per subject.

## Discussion

In this study, we used cortical tracking of continuous speech, in a multi-talker scenario, to investigate the neural representations of an auditory scene. From MEG recordings of subjects selectively attending to one of the three co-located speech streams, we observed that 1) The early neural responses (from sources with short latencies), which originate primarily from core auditory cortex, represent the foreground (attended) and background (ignored) speech streams without any significant difference, whereas the late neural responses (from sources with longer latencies), which originate primarily from higher-order areas of auditory cortex, represent the foreground with significantly higher fidelity than the background; 2) Early neural responses are not only balanced in how they represent the constituent speech streams, but in fact represent the entire acoustic scene holistically, rather than as separately contributing individual perceptual objects; 3) Even though there are two physical speech streams in the background, no neural segregation is observed for the background speech streams.

It is well established that auditory processing in cortex is performed in a hierarchical fashion, in which an auditory stimulus is processed by different anatomical areas at different latencies (Inui et al., 2006; Nourski et al., 2014). Using this idea to inform the neural decoding/encoding analysis allows the effective isolation of neural signals from a particular cortical area, and thereby the ability to track changes in neural representations as the stimulus processing proceeds along the auditory hierarchy. This time-constrained reconstruction/prediction approach may prove especially fruitful in high-time-resolution/low-spatial-resolution imaging techniques such as MEG and EEG. Even though different response components are generated by different neural sources, standard neural source localization algorithms may perform poorly when different sources are strongly correlated in their responses (Lutkenhoner and Mosher, 2007). While the proposed method is not to be viewed as an alternative to source localization methods, it can nonetheless be used to tease apart different components of MEG/EEG response, without explicit source localization.

Even though there is no significant difference between the ability to reconstruct the foreground and background from early neural responses, nonetheless we observe a non-significant tendency towards an enhanced representation of the foreground (foreground > background, *p* = 0.21). This could be due to task-related plasticity of spectro-temporal receptive fields of neurons in mammalian primary auditory cortex (Fritz et al., 2003), where the receptive fields of neurons are tuned to match the stimulus characteristics of attended sounds. The selective amplification of foreground in late neural responses (from higher-order auditory cortices) but not in early responses (from core auditory cortex) observed here using *decoding* is in agreement with the *encoding* result of Ding and Simon (2012b) where the authors showed that the late M100_TRF_ component, but not the early M50_TRF_ component, of TRF is significantly modulated by attention. The increase in fidelity of the foreground as the response latency increases indicates a temporal as well as functional hierarchy in cortical processing of auditory scene, from core to higher-order areas in auditory cortex. Similar preferential representation for the attended speech stream has been demonstrated, albeit with only two speech streams and not differentiating between early and late responses, using delta and theta band neural responses (Ding and Simon, 2012b; Zion Golumbic et al., 2013a; Zion Golumbic et al., 2013b) as well as high-gamma neural responses (Mesgarani and Chang, 2012; Zion Golumbic et al., 2013a), and using monaural (Ding and Simon, 2012b; Mesgarani and Chang, 2012) as well as audio-visual speech (Zion Golumbic et al., 2013a; Zion Golumbic et al., 2013b).

While some researchers suggest selective entrainment (Schroeder and Lakatos, 2009; Ng et al., 2012; Zion Golumbic et al., 2013b; Kayser et al., 2015) as the mechanism for selective tracking of attended speech, others suggest a temporal coherence model (Shamma et al., 2011; Ding and Simon, 2012b). Natural speech is quasi-rhythmic with dominant rates at syllabic, word and prosodic frequencies. The selective entrainment model suggests that attention causes endogenous low frequency neural oscillations to align with the temporal structure of the attended speech stream, thus aligning the high excitability phases of oscillations with events in attended stream. This effectively forms a mask that favors the attended speech. The temporal coherence model suggests that selective tracking of attended speech is achieved in two stages. First, a cortical filtering stage, where feature-selective neurons filter the stimulus, producing a multidimensional representation of auditory scene along different feature axes. This is followed by a second stage, coherence analysis, which combines relevant features streams based on their temporal similarity, giving rise to separate perceptions of attended and ignored streams. In this model, it is hypothesized that attention, acting through in the coherence analysis stage, plays an important role in stream formation. This type of coherence model predicts an unsegregated representation of any (non-attended) background streams.

The representation of an auditory scene in core auditory cortex, based on the early responses, is here shown to be more spectro-temporal- or acoustic-based than object-based (e.g., Figure 3B). This is further supported by the result that the Early-late model predicts MEG neural responses significantly better than Summation model (e.g., Figure 5). This is consistent with previous demonstrations that neural activity in core auditory cortex was highly sensitive to acoustic characteristics of speech and primarily reflects spectro-temporal attributes of sound (Nourski et al., 2009; Okada et al., 2010; Ding and Simon, 2013; Steinschneider et al., 2014). In contrast, Nelken and Bar-Yosef (2008) suggest that neural auditory objects may form as early as primary auditory cortex, and Fritz et al. (2003) show that representations of dynamic sounds in primary auditory cortex are influenced by task. As a working principle, it is possible that less complex stimuli are resolved earlier in the hierarchy of auditory pathway (e.g., sounds that can be separated via tonotopy) whereas more complex stimuli (e.g., concurrent speech streams), which need further processing, are resolved only much later in auditory pathway. In addition, it is worth noting that the current study uses co-located speech streams, whereas mechanisms of stream segregation will also be influenced by other auditory cues, including spatial cues, differences in acoustic source statistics (e.g., only speech streams vs. mixed speech and music; strong statistical differences might drive stream segregation in a more bottom-up manner than the top-down attentional effects studied here), perceptual load effects (e.g., tone streams vs. speech streams), as well as visual cues. Any of these additional cues has the potential to alter the timing and neural mechanisms by which auditory scene analysis occurs.

It is widely accepted that an auditory scene is *perceived* in terms of auditory objects (Bregman, 1994; Griffiths and Warren, 2004; Shinn-Cunningham, 2008; Shamma et al., 2011). Ding and Simon (2012a) demonstrated evidence for an object-based cortical representation of an auditory scene, but did not distinguish between early and late neural responses. This, coupled with the result here that early neural responses provide an acoustic, not object-based, representation, strongly suggest that the object-based representation emerges only in the late neural responses/higher-order (belt and parabelt) auditory areas. This is further supported by the observation that acoustic invariance, a property of object-based representation, is observed in higher order areas but not in core auditory cortex (Chang et al., 2010; Okada et al., 2010). When the foreground is represented as an auditory object in late neural responses, the finding that the combined background is better reconstructed than the sum of envelopes of individual backgrounds (Figure 4B) suggests that in late neural responses the background is not represented as separated and distinct auditory objects. This result is consistent with that of Sussman et al. (2005), who reported an unsegregated background when subjects attended to one of three tone streams in the auditory scene. This unsegregated background may be a result of an ‘analysis-by-synthesis’ (Yuille and Kersten, 2006; Poeppel et al., 2008) mechanism, wherein the auditory scene is first decomposed into basic acoustic elements, followed by top-down processes that guide the synthesis of the relevant components into a single stream, which then becomes the object of attention. The remainder of the auditory scene would be the unsegregated background, which itself might have the properties of an auditory object. When attention shifts, new auditory objects are correspondingly formed, with the old ones now contributing to the unstructured background. Shamma et al. (2011) suggest that this top down influence acts through the principle of temporal coherence. Between the two opposing views, that streams are formed pre-attentively and that multiple streams can co-exist simultaneously, or that attention is required to form a stream and only that single stream is ever present as separated perceptual entity, these findings lend support to the latter.

In summary, these results provide evidence that, in a complex auditory scene with multiple overlapping spectral and temporal sources, the core areas of auditory cortex maintains an acoustic representation of the auditory scene with no significant preference to attended over ignored source, and with no separation into distinct sources. It is only the higher-order auditory areas that provide an object based representation for the foreground, but even there the background remains unsegregated.

